# CoCoNat: a novel method based on deep-learning for coiled-coil prediction

**DOI:** 10.1101/2023.05.08.539816

**Authors:** Giovanni Madeo, Castrense Savojardo, Matteo Manfredi, Pier Luigi Martelli, Rita Casadio

## Abstract

**Motivation:** Coiled-coil domains (CCD) are widespread in all organisms performing several crucial functions. Given their relevance, the computational detection of coiled-coil domains is very important for protein functional annotation. State-of-the art prediction methods include the precise identification of coiled-coil domain boundaries, the annotation of the typical heptad repeat pattern along the coiled-coil helices as well as the prediction of the oligomerization state.

**Results:** In this paper we describe CoCoNat, a novel method for predicting coiled-coil helix boundaries, residue-level register annotation and oligomerization state. Our method encodes sequences with the combination of two state-of-the-art protein language models and implements a three-step deep learning procedure concatenated with a Grammatical-Restrained Hidden Conditional Random Field (GRHCRF) for CCD identification and refinement. A final neural network (NN) predicts the oligomerization state. When tested on a blind test set routinely adopted, CoCoNat obtains a performance superior to the current state-of-the-art both for residue-level and segment-level coiled-coil detection. CoCoNat significantly outperforms the most recent state-of-the art method on register annotation and prediction of oligomerization states.

**Availability:** CoCoNat is available at https://coconat.biocomp.unibo.it.

## 1 Introduction

Coiled-coil domains (CCD) in proteins are structural motifs where α-helices pack together in an arrangement called knobs-into-holes (Crick 1952, Crick, 1953a, 1953b). Since the first crystallographic observation in the structure of influenza virus hemagglutinin (Wilson *et al*., 1981), CCDs have been resolved in several proteins through all the kingdoms of life (Truebestein and Leonard, 2016). CCDs are present, among others, in structural proteins, transcription factors, and enzymes (Truebestein and Leonard, 2016; Lupas and Bassler, 2017). CCDs act as molecular spacers, influence the organelle organization, constrain the distance of residues involved in binding and catalytic sites, mediate membrane fusion, and are involved in signal transduction and solute transport.

Canonical CCDs include the interaction of two, three or four α-helices. Each helix is characterized by the repetition of a seven-residue motif (heptad repeat) whose positions are referred to as registers and are labelled as *abcdefg*. Positions *a* and *d* are routinely occupied by hydrophobic residues and mediate the interaction between different helices in the domain. Since the periodicity of α-helices in CCDs is only 3.5 residues per turn (in contrast with canonical α-helices, in which the periodicity is 3.6), residues in the same register lie on the same side of the helix surface. Therefore, the hydrophobic nature of residues in registers *a* and *d* confers a peculiar amphipathic character to the α-helix. In CCDs, α-helices interact with each other through their hydrophobic face (Lupas and Gruber, 2005). Rarely, CCDs contain helices characterized by non-canonical repeats, longer than 7 residues, i.e.: hendecades, pentadecades and nonadecades (Lupas and Gruber, 2005; Szczepaniak *et al*., 2020). This paper and other studies in the field do not include these cases.

CCDs are classified according to the number and orientation of the involved α-helices, i.e., by their oligomerization state. CCDs based on the orientation of helices are classified as parallel or antiparallel and based on the number of helices as dimers, trimers, tetramers.

CCDs are routinely annotated starting from the protein 3D structure, adopting specialized software such as SOCKET (Walshaw and Woolfson, 2001) and SamCC-Turbo (Szczepaniak *et al*., 2020). Annotations performed with the two methods are collected in the CC+ database (http://coiledcoils.chm.bris.ac.uk/ccplus/search/, Testa *et al*., 2009) and in the CCdb database (https://lbs.cent.uw.edu.pl/ccdb), respectively. Semimanual annotations are also available in the SCOPe database (Fox *et al*., 2014).

The relevance of CCDs in protein annotation requires the development of computational methods for predicting the presence and localization of CCDs (including registers), and their oligomerization state, starting from the protein sequence. Over the years, several methods have been proposed, addressing the different tasks of CCD prediction. Boundaries of α-helices involved in CCDs can be predicted with COILS (Lupas *et al*., 1991), PCOILS (Gruber *et al*., 2005), MarCoil (Delorenzi and Speed, 2002), MultiCoil2 (Trigg *et al*., 2011), CCHMM_PROF (Bartoli *et al*., 2009), DeepCoil (Ludwiczak *et al*., 2019), and CoCoPred (Feng *et al*., 2022).The oligomeric state is predicted with PrOCoil (Mahrenholz *et al*., 2011), LOGICOIL (Vincent *et al*., 2013), MultiCoil2 (Trigg *et al*., 2011) and CoCoPred (Feng *et al*., 2022). To date, only CoCoPred predicts the registers along the heptads (Feng *et al*., 2022).

Recently protein language models improved sequence encoding procedures. Here we introduce CoCoNat which, for the first time, adopts a sequence encoding, based on the combination of two state-of-the-art protein language models, ProtT5 (Elnaggar *et al*., 2021) and ESM1-b (Rives *et al*., 2021) and based on deep-learning computes: i) the coiled-coil helix boundaries; ii) the residue-level register annotation, and iii) the CCD oligomerization state.

We trained CoCoNat on a dataset comprising 2198 proteins containing CCDs and 9062 proteins without CCD (negative examples). When tested on a blind test set including 439 CCD and 279 non-CCD proteins, CoCoNat scores with a performance that is superior to the current state-of-the-art both for residue-level and segment-level CCD detection. Moreover, CoCoNat significantly outperforms the most recent method, CoCoPRED (Feng *et al*., 2022), on both register annotation as well as prediction of CCD oligomerization states.

CoCoNat is available as web server at https://coconat.biocomp.unibo.it.

## 2 Methods

### 2.1 Datasets

CoCoNat is trained and tested on the same datasets adopted in CoCoPRED (Feng *et al*., 2022). Numbers are summarized in Table 1 and all datasets are available at the CoCoNat website (https://coconat.biocomp.unibo.it).

**Table 1.** Training and testing set of CoCoNat

|  | Positive Proteins | Helices in CCDs | Negative Proteins |
| --- | --- | --- | --- |
| **Training Set** | 2191 | 4342 | 9040 |
| **Blind test Set** | 439 | 937 | 279 |

#### 2.1.1 Training dataset

The positive training dataset contains 2191 proteins out of the 2337 included in CoCoPRED and deriving from 30,227 CCD-containing proteins annotated with SOCKET (Walshaw and Woolfson, 2001) in the CC+ database (Testa *et al*., 2009).

Briefly, proteins included in the positive training set of CoCoPRED have been selected with the following criteria: i) protein structure resolution < 4 Å, ii) protein length between 25 and 700 residues, iii) length of CCD helices ≥ 8 residues; iv) absence of non-canonical (i.e., non-heptad based) CCDs; v) CCD oligomeric state classified as parallel or antiparallel dimer, trimer, and tetramer; vi) sequence pairwise identity lower than 30% with respect to proteins in the testing set (see below); vii) internal pairwise sequence identity lower than 50%.

Since CoCoNat relies on full length proteins for the computation of sequence embeddings, we applied further filters to the CoCoPRED positive training dataset, removing: i) proteins not mapped into UniProt; ii) synthetic and fusion proteins; iii) proteins whose structure coverage with respect to the UniProt sequence is lower than 70%. After this screening, the positive dataset includes 2191 proteins with 4342 coiled-coil helices, whose length ranges from 8 to 145 residues. The number of coiled-coil helices per protein ranges from 1 to 19.

The negative training set of CoCoNat includes 9040 proteins. This derives from the 9358 proteins of the negative set CoCoPRED that was obtained from the negative set of DeepCoil (Ludwiczak *et al*., 2019) after the exclusion of proteins with a sequence identity>30% with respect to the blind test set (see below) and with a sequence identity>50% with respect to the positive training set. We filtered out proteins not mapped into UniProt.

Proteins in the training set (both positive and negative examples) were split into five subsets for 5-fold cross-validation. To reduce the redundancy among cross-validation sets, proteins sharing more than 25% sequence identity at 50% coverage are clustered in the same set. Cross-validation sets were used to set all the hyperparameters.

#### 2.1.2 Blind test dataset

To test and benchmark CoCoNat with other available methods, we adopted the 718 proteins (439 with CCDs and 279 without CCDs) included in the CoCoPRED test set. The CoCoPRED set shares less than 30% sequence identity with proteins in the training sets of CCHMM_PROF (Bartoli *et al*., 2009), MARCOIL (Delorenzi and Speed, 2002), and Multicoil2 (Trigg et al., 2011), CoCoPRED (Feng *et al*., 2022) and DeepCoil (Ludwiczak *et al*, 2019).

Following the annotation of CoCoPred, the 439 CCDs include 937 coiled-coil helices, whose length ranges between 8 and 71 residues. The number of coiled-coil helices per protein is from 1 to 8.

For the annotation of registers and oligomeric states we run SOCKET in house and obtained coherent annotations for 429 of the 439 proteins with CCDs, including 808 coiled-coil helices.

### 2.2 Protein encoding

CoCoNat makes use of residue embeddings obtained with large-scale protein Language Models (pLMs) to represent proteins in training and testing sets. Specifically, we adopted two state-of-the-art pLMs: ProtT5 (Elnaggar *et al*., 2021) and ESM-1b (Rives *et al*., 2021), generating, for each residue in the protein sequence, 1024 and 1280 features, respectively. Residue-level representations are then concatenated together, leading to vectors of 2304 dimensions for each residue in the sequences. The concatenation of embeddings obtained with different pLMs has been shown to improve the performance in previous works (Manfredi *et al*., 2022, 2023).

### 2.3 CoCoNat architecture

CoCoNat is organized as a three-step method combining a deep learning approach, a probabilistic graphical model, and a single-layer neural network in a cascading way. The first two steps are collectively devised to detecting coiled-coil helix boundaries and the residue-level annotation of registers within each predicted helix. The third step predicts the CCD oligomerization state (Figure1). In the following, each step is briefly described.

**Figure 1.**
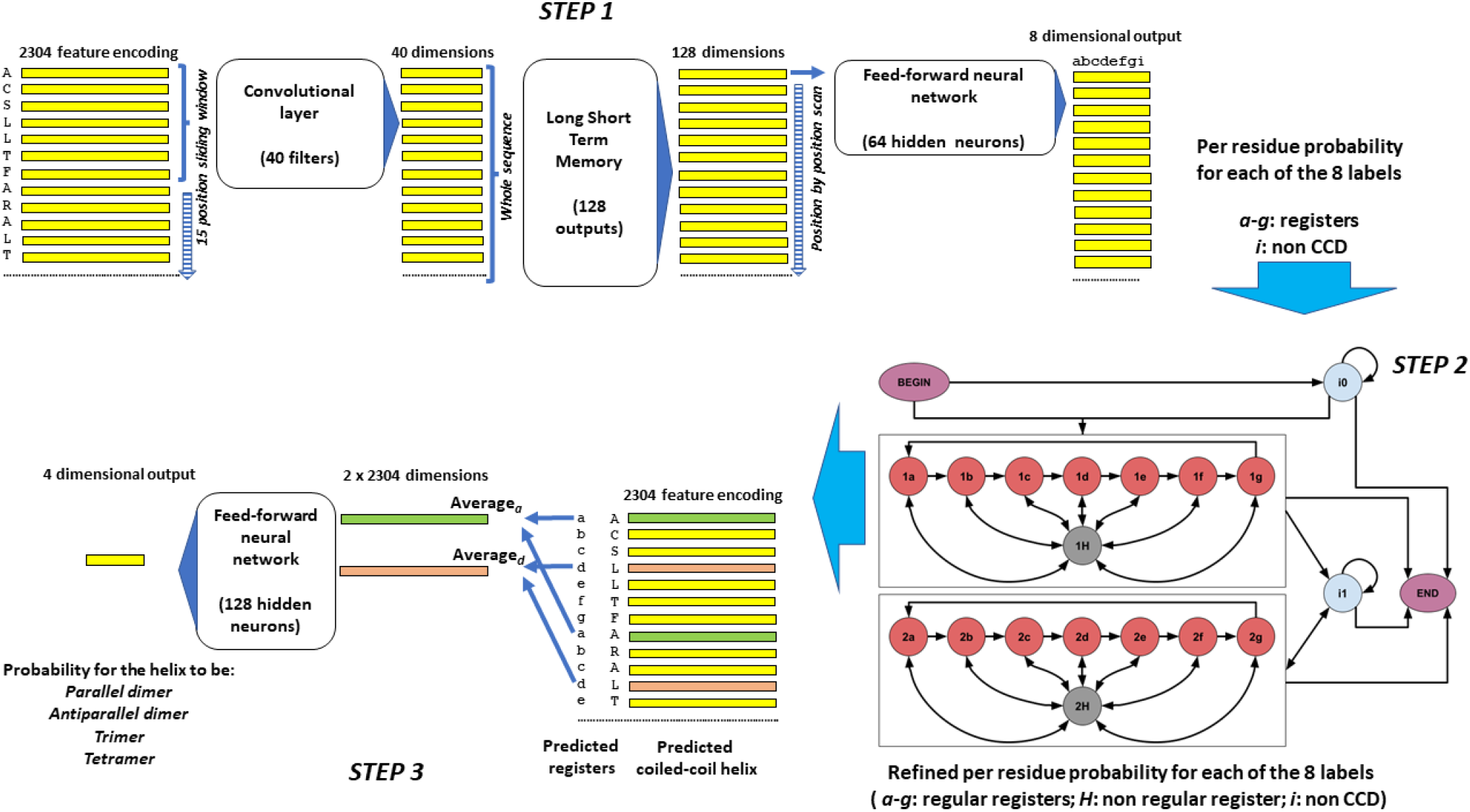
Workflow of the predictive model of CoCoNat, comprising three steps. The *STEP 1* maps each residue, encoded with a 2304 dimensional vector (the concatenation of ProtT5 and ESM1-b embeddings) into an 8-dimensional vector representing the probability distribution over the labels (*a-g* registers for coiled-coil helices and *i* for non-CCD portions). It combines in cascade i) a convolutional layer that computes a 40-dimensional representation for each position in the sequence, based on a sliding window of 15 contiguous residues, ii) a Long Short-Term Memory layer that analyzes the whole sequence and provides a 128-dimensional representation for each position, and iii) a fully connected Feed-Forward Neural Network that, residue by residue, provides the 8-dimensional mapping. The *STEP 2* consists of a Grammatical Restrained Hidden Conditional Random Field (GRHCRF) that casts the grammar of CCDs in the topology of the connections among 20 different states. Each sequence is generated by a path that starts from the BEGIN state and can either enter the self-looping *i0* state, which models the N-terminal non-CCD portion of the protein, or the first 8-state block (*1a-1b-1c-1d-1e-1f-1g-1H*), which model the first CCD domain. Labels *a-g* of the states in the block correspond to registers and state *H* accommodates non regular transitions. Residues after the first coiled-coil helix are modelled by the *i1* state (non-CCD) and, in case, by a second CCD block (*2a-2b-2c-2d-2e-2f-2g-2H*), analogous to the first one. All states, but *1H* and *2H*, can make transition to the END state, terminating the path. GRHCRF provides the annotation of coiled-coil helix boundaries and of registers by computing the optimal a-posteriori Viterbi path, given the probabilities computed in the STEP 1. The *STEP 3* provides the prediction of the oligomeric state, based on the annotation computed in STEP 2 and on the 2304 -dimensional embedding. For each predicted coiled-coil helix, embeddings labelled with registers *a* and *d* are separately averaged and fed into a Feed-Forward Neural Network that computes the probability distribution over the four possible classes (parallel and antiparallel dimer, trimer, and tetramer).

#### 2.3.1 Deep learning architecture

The first step (Figure 1) is based on one convolutional layer (LeCun *et al*., 1989) followed by a Long Short-Term Memory (LSTM) layer (Hochreiter and Schmidhuber, 1997). The convolutional layer captures local depend-encies of the input data. This layer, adopting a 15 residue long sliding window, takes as input the protein, where each residue is represented with a 2304-feature vector. By applying 40 different filters, the layer outputs the same protein with residues encoded with 40-feature vectors. This mapping is provided as input to a LSTM layer including 128 output neurons, which captures long-range dependencies. Finally, we apply a standard feed-forward network (with 64 hidden neurons) with 8 output neurons (one for each possible coiled-coil registers, *a*-*b-c-d-e-f-g*, plus one (*i*) for non-CCD residues), endowed with a sigmoid activation function. The output gives the per-residue probability of each register or none.

In order to reduce overfitting, we introduce dropout layers between convolutional, LSTM, and the feed forward network with rate fixed to 0.25.

The architecture was trained by adopting a five-fold cross validation, with a gradient descent computed by the Kullback-Leibler divergence error function and the Adam optimization algorithm (Kingma and Ba, 2017). The best model was determined using the early stopping technique of 10 epochs in which the validation error did not decrease. To implement the architecture, we use the Keras Python library (Chollet, 2015). The method hyperparameters have been optimized during cross validation.

#### 2.3.2 Refining the prediction with Grammatical-Restrained Hidden Conditional Random Field (second step)

The second step (Figure 1) inputs protein mapping as computed by step 1. GRHCRF is a discriminative probabilistic model (Fariselli *et al*., 2009, Madeo et al., 2021), and it allows to introduce the regular grammar of the CDD registers. In the prediction phase, a Posterior-Viterbi dynamic-programming algorithm computes the most probable path along the model, satisfying the grammatical constraints.

In our model, the grammar has two identical blocks for CCD prediction, two states (i0 and i1 in Figure 1, step 2) with a self-loop modeling the non-CCD regions and one BEGIN and one END states. In each CCD block, seven states model the register sequence, and one further state (H, 1 and 2) accommodates non-regular transitions (i.e., transitions escaping the regular heptad repeat pattern, *abcdefg*). The first GRHCRF block models the first CC helix and the second one models all the others CC helices, when present, in the protein sequence.

The GRHCRF output defines the precise identification of coiled-coil domain boundaries and annotates the typical heptad repeat pattern along the coiled-coil helices.

#### 2.3.3 Prediction of the oligomerization state

The prediction of the CCD oligomerization state adopts a simple feed-forward neural network with a single hidden layer, comprising 128 neurons, and four output units corresponding to the four possible oligomerization states: parallel and antiparallel dimers, trimers, and tetramers.

The input of this network is built based on a well-known biophysical feature: the oligomeric state of canonical CCD is largely determined by the nature of hydrophobic residues in the heptad repeat pattern, namely, residues labelled with registers *a* and *d* (see Woolfson, 2023). Based on this observation, for a given coiled-coil helix, the network input is obtained concatenating the average embedding vectors of *a* and *d* predicted positions.

More formally, given a coiled-coil helix of length *l*, with an embedding matrix *E* of dimension *l* × 2304 (as derived from the concatenation of two pLM embeddings, ProtT5 and ESM1-b), the following two mean vectors are computed:

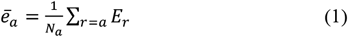

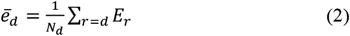

where *Na* and *Nd* are the number of positions labeled with registers *a* and *d*, respectively. The input vector for the network is then obtained con-catenating ē_a_ and ē_d_:

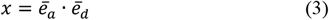

where [·] denotes the vector concatenation operator.

The NN has been implemented with the Python PyTorch package (https://pytorch.org/), selecting a categorical cross-entropy error function and the Adam optimizer.

### 2.4 Scoring performance

To evaluate the performance of our method in recognizing coiled-coil helices, we adopted residue- and segment-based measures.

The residue-based scores include precision (PRER), recall (RECR), and F1-score (F1R).

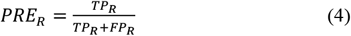

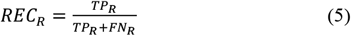

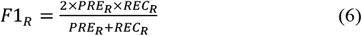

where TPR, FPR, and FNR are true positive, false positive, and false negative coiled-coiled residues, respectively.

Analogously, the segment-based scores include precision (PRES), recall (RECS), and F1-score (F1S):

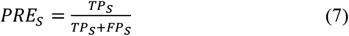

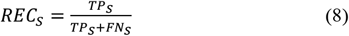

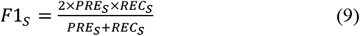

In this case, TPS, FPS and FNS are computed for the coiled-coil helices. Prediction is considered correct (TPS) only if the overlap between predicted and observed segments is at least equal to the half-length of the longest segment.

Following Feng *et al*. (2022), we used two Segment Overlap (SOV) measures, one taking as reference observed residues (SOVo) and one taking as reference predicted residues (SOVp) (Zemla *et al*., 1999).

We computed the Precision-Recall (PR) curve and the relative Area Under the Curve (PR-AUC), by plotting the two measures at varying thresholds of coiled-coil probabilities as obtained from the GRHCRF posterior probability values.

For the register and oligomeric state prediction tasks, we reported our results as confusion matrices. Additionally, we computed class level MCC values.

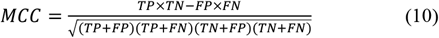

## 3 Results

### 3.1 Cross-validation results

We performed 5-fold cross-validation experiments to optimize the input encoding of our model. To evaluate the contribution from the two protein language models, we independently trained three different identical models adopting as input: i) ProtT5, ii) ESM1-b, and iii) both encodings combined in a single vector. Results are reported in Table 2. The overall prediction achieved with each one of the two language models is similar (F1R and F1S scores equals 0.44, 0.34 and 0.45, 0.33 with ProtT5 and ESM1-b, respectively). However, the ProtT5-based input produces more sensitive than precise results (e.g., residue-level precision and recall are 0.42 and 0.46, respectively), whereas input based on ESM1-b provides more precise than sensitive predictions (residue-level precision and recall are 0.49 and 0.41, respectively). Combining the two embeddings into a single vector leads to better performances, raising both precision and recall value, and achieving F1R and F1s values of 0.49 and 0.37, respectively. These results agree with previous works in which the combination of embeddings from different pLMs has been proven effective also for other prediction tasks (Manfredi *et al*., 2022, 2023). All results presented in this manuscript are obtained using the combination of the two input embeddings.

**Table 2.** Prediction of coiled-coil helices with CoCoNat adopting different embeddings.

| Input | PRE<sub>R</sub> | REC<sub>R</sub> | F1<sub>R</sub> | PRE<sub>S</sub> | REC<sub>S</sub> | F1<sub>S</sub> |
| --- | --- | --- | --- | --- | --- | --- |
| ProtT5 | 0.42 | 0.46 | 0.44 | 0.32 | 0.36 | 0.34 |
| ESM | 0.49 | 0.41 | 0.45 | 0.34 | 0.33 | 0.33 |
| ProtT5+ESM | 0.51 | 0.47 | 0.49 | 0.38 | 0.37 | 0.37 |

### 3.2 Prediction of coiled coils on the blind test set

CoCoNat is benchmarked on the same blind test set adopted by CoCoPRED, which was in turn compared to pre-existing available methods (Feng *et al*., 2022). In Table 3 we report our results together with those already published by Feng *et al*. (2022), for both at residue- and segment-level predictions. The performances of CoCoNat, which adopts encodings based on ProT5 and ESM1-b well compare with state-of-the-art, showing (with respect to the top performing methods in the benchmark) an improvement in the per-residue precision value (0.48), with a slight loss in recall (0.57), which is reflected in the higher value of the F1-score (0.52). The per-segment scores of CoCoNat confirm this trend. Moreover, CoCoNat performs with the highest SOVp and the second highest SOVo (see Section 2.4 for definition).

**Table 3.** CoCoNat and the state-of-art method on the same blind test set

| Method* | PRE<sub>R</sub> | REC<sub>R</sub> | F1<sub>R</sub> | PRAUC | PRE<sub>S</sub> | REC<sub>S</sub> | F1<sub>S</sub> | SOV<sub>O</sub> | SOV<sub>P</sub> |
| --- | --- | --- | --- | --- | --- | --- | --- | --- | --- |
| COILS | 0.34 | 0.27 | 0.3 | 0.26 | 0.22 | 0.06 | 0.01 | 18.92 | 39.95 |
| MarCoil | 0.28 | 0.34 | 0.3 | 0.25 | 0.15 | 0.07 | 0.09 | 22.53 | 33.73 |
| PCOILS | 0.3 | 0.34 | 0.32 | 0.27 | 0.16 | 0.07 | 0.09 | 22.73 | 35.72 |
| CCHMM_prof | 0.14 | **0.63** | 0.23 | - | 0.11 | 0.26 | 0.15 | 44.34 | 17.51 |
| MultiCoil2 | 0.33 | 0.22 | 0.27 | 0.24 | 0.14 | 0.01 | 0.03 | 10.49 | 26.19 |
| DeepCoil2 | 0.42 | 0.61 | 0.5 | 0.47 | 0.29 | 0.47 | 0.36 | 53.72 | 49.49 |
| CoCoPRED | 0.43 | 0.61 | 0.5 | **0.5** | 0.42 | **0.54** | 0.47 | **61.91** | 50.28 |
| CoCoNat | **0.48** | 0.57 | **0.52** | 0.47 | **0.48** | 0.51 | **0.49** | 60.73 | **57.71** |

\*COILS (Lupas *et al.*, 1991); MarColi (Delorenzi and Speed, 2002); PCOILS (Gruber *et al.*, 2005); CCHMM\_prof (Bartoli *et al.*, 2009); MultiCoil2 (Trigg *et al.*, 2011); DeepCoil2 (Ludwiczak *et al.*, 2019); CoCoPRED (Feng *et al.*, 2022). Subscript R: per residue; subscript S: per CC helix segment. Results are derived from Feng *et al.* (2022), with the exception of CoCoNat. The blind test set contains 439 positive and 279 negative proteins (Feng *et al.*, 2022).

### 3.3 Prediction of coiled-coil registers

We compared CoCoNat and CoCoPRED in the task of annotating heptad repeat registers. We used the blind test of 429 proteins endowed with SOCKET annotations to ensure a fair comparison between the two approaches. Results of CoCoPRED were generated using the standalone version of the program (Figure 2).

**Figure 2.**
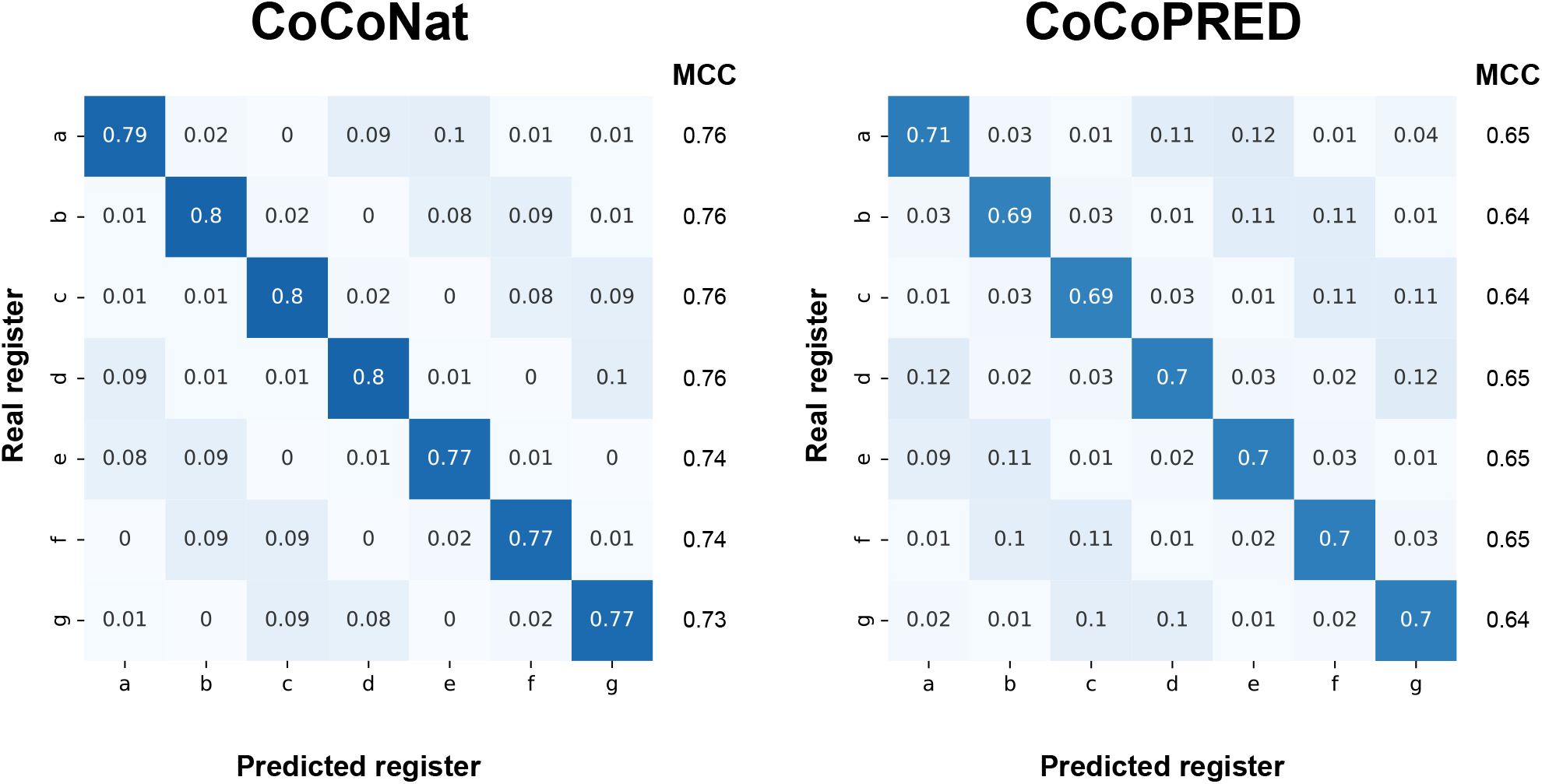
Confusion matrices (represented as heatmaps) and class-level MCC scores for register prediction obtained on the blind test set with CoCoNat (left) and CoCoPRED (right). The results refer to 429 test proteins endowed with CCDs for which SOCKET assigned a coherent annotation. Values on the main diagonal of the matrix estimate the correct recall rate in each class.

CoCoNat outperforms CoCoPRED. For all the register labels *a-g*, CoCoNat recall values (reported in the diagonal of the confusion matrix in Figure 3) are higher by 7-11 percentage points with respect to corresponding CoCoPRED scores. Furthermore, MCC values indicate an improvement ranging from 7 to 12%.

**Figure 3.**
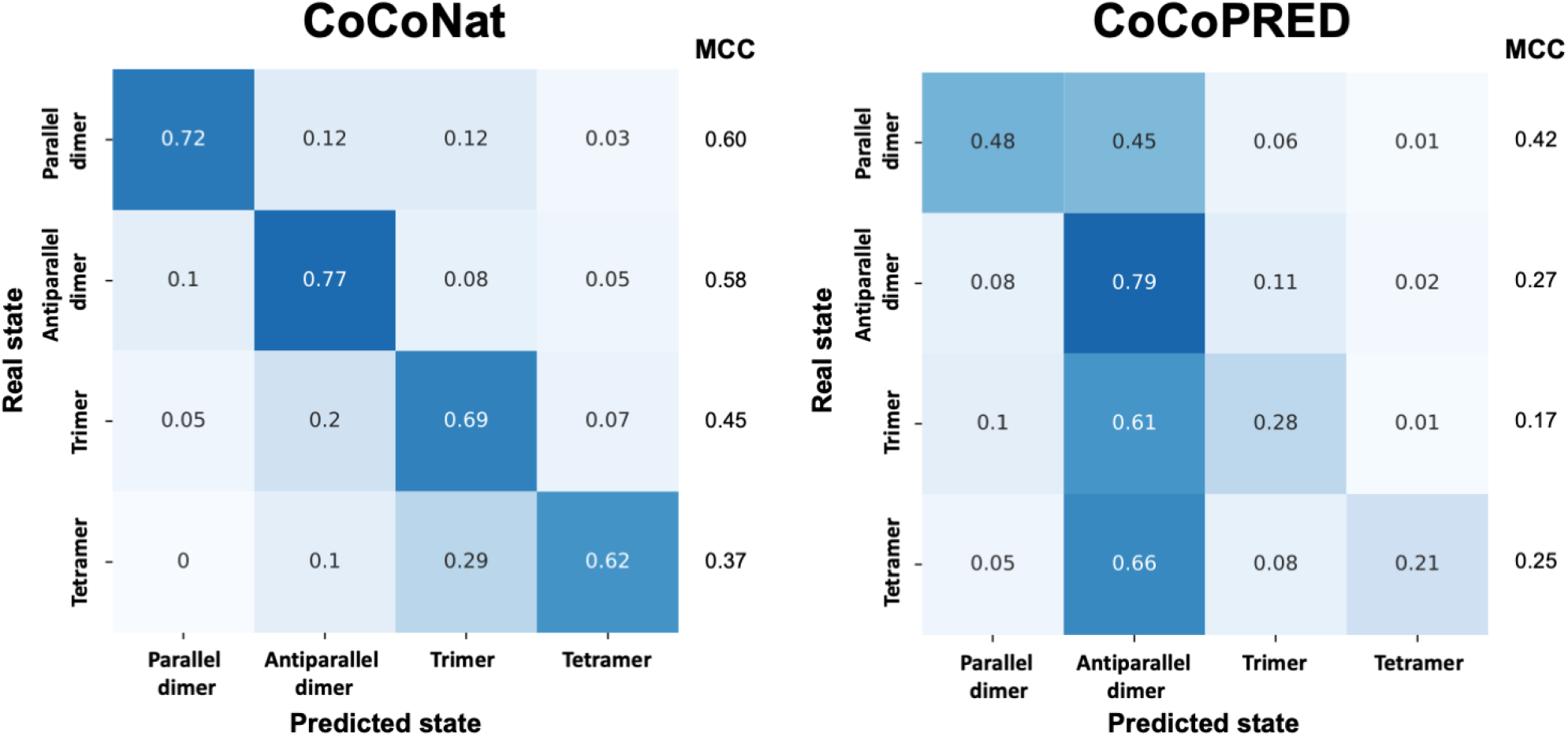
Confusion matrices (represented as heatmaps) and class-level MCC scores for the oligomeric state prediction obtained on the blind test set with CoCoNat (left) and CoCoPRED (right). The results refer to 429 test proteins endowed with CCDs for which SOCKET provided an annotation. Values on the main diagonal of the matrix estimate the correct recall rate in each class.

Remarkably, CoCoNat recall and MCC values are quite similar across all the 7 different register labels, suggesting that the register is routinely predicted in the correct regular configuration, from label *a* to *g*. This highlights that the GRHCRF grammar is properly capturing transition constraints among the different registers inside the coiled-coil segments.

Wrong CoCoNat assignments (about 10%) mostly depend on an erroneous shift among *a* and *d* registers.

### 3.4 Prediction of the CCD oligomerization state

We finally compared CoCoNat and CoCoPRED on the task of predicting CCD oligomerization state. Again, we used the blind test set of 429 proteins as benchmark. The two methods are compared assuming an oracle predictor for the identification of CCD segments (i.e., we classify real CCD segments into the four oligomerization classes). In Figure 3 we report a side-by-side comparison of the two approaches in terms of confusion matrices.

Looking at the recall values reported in the diagonal of the two confusion matrices, CoCoNat significantly overpasses CoCoPRED, providing predictions that are overall more balanced across the four oligomerization state classes. Remarkably, CoCoNat outperforms CoCoPRED also on less abundant classes i.e., trimers and tetramers, as it is evident from the comparison of class-level MCC values for parallel dimers, antiparallel dimers, trimers, and tetramers.

## 4 Conclusion

In this paper we described CoCoNat, a novel method based on protein language model embeddings and deep learning for detection of coiled-coiled helices at residue level, prediction of coiled-coil heptad repeat registers and oligomerization state.

Training and testing were performed on datasets derived from literature. When compared with other state-of-the-art tools, CoCoNat reported performance that are significantly better than those obtained by other approaches tested, in particular, when considering register and oligomerization state prediction.

In this work we also proved the relevance of adopting protein residue representations derived from large-scale protein language models such as ProtT5 (Elnaggar *et al*., 2021) and ESM1-b (Rives *et al*., 2021) for this specific task. Moreover, we further confirmed that the combination of different language models provides better performance, suggesting that different models obtained with different architectures and data give complementary representations.

We release CoCoNat as web server available at https://coconat.bio-comp.unibo.it, from which the user can analyze single protein as well as larger datasets containing up to 500 sequences per job.

## References

Bartoli, L. et al. (2009) CCHMM_PROF: a HMM-based coiled-coil predictor with evolutionary information. Bioinformatics, 25, 2757–2763.

Chollet, F. (2015) Keras GitHub.

Crick, F.H.C. (1952) Is alpha-keratin a coiled coil? Nature, 170, 882–883.

Crick, F.H.C. (1953a) The Fourier transform of a coiled-coil. Acta Cryst, 6, 685–689.

Crick, F.H.C. (1953b) The packing of α-helices: simple coiled-coils. Acta Cryst, 6, 689–697.

Delorenzi, M. and Speed, T. (2002) An HMM model for coiled-coil domains and a comparison with PSSM-based predictions. Bioinformatics, 18, 617–625.

Elnaggar, A. et al. (2021) ProtTrans: Towards Cracking the Language of Lifes Code Through Self-Supervised Deep Learning and High Performance Computing. IEEE Trans. Pattern Anal. Mach. Intell., 1–1.

Fariselli, P. et al. (2009) Grammatical-Restrained Hidden Conditional Random Fields for Bioinformatics applications. Algorithms Mol Biol, 4, 13.

Feng, S.-H. et al. (2022) CoCoPRED: coiled-coil protein structural feature prediction from amino acid sequence using deep neural networks. Bioinformatics, 38, 720–729.

Fox, N.K. et al. (2014) SCOPe: Structural Classification of Proteins—extended, integrating SCOP and ASTRAL data and classification of new structures. Nucl. Acids Res., 42, D304–D309.

Gruber, M. et al. (2005) REPPER--repeats and their periodicities in fibrous proteins. Nucleic Acids Res, 33, W239-243.

Hochreiter, S. and Schmidhuber, J. (1997) Long Short-Term Memory. Neural Computation, 9, 1735–1780.

Kingma, D.P. and Ba, J. (2017) Adam: A Method for Stochastic Optimization. arXiv:1412.6980 [cs].

LeCun, Y. et al. (1989) Backpropagation Applied to Handwritten Zip Code Recognition. Neural Computation, 1, 541–551.

Li, C. et al. (2016) Critical evaluation of in silico methods for prediction of coiled-coil domains in proteins. Brief Bioinform, 17, 270–282.

Ludwiczak, J. et al. (2019) DeepCoil-a fast and accurate prediction of coiled-coil domains in protein sequences. Bioinformatics, 35, 2790–2795.

Lupas, A. et al. (1991) Predicting coiled coils from protein sequences. Science, 252, 1162–1164.

Lupas, A.N. and Bassler, J. (2017) Coiled Coils - A Model System for the 21st Century. Trends Biochem Sci, 42, 130–140.

Lupas, A.N. and Gruber, M. (2005) The Structure of α-Helical Coiled Coils. In, Advances in Protein Chemistry, Fibrous Proteins: Coiled-Coils, Collagen and Elastomers. Academic Press, pp. 37–38.

Madeo, G. et al. (2021) BetAware-Deep: An Accurate Web Server for Discrimination and Topology Prediction of Prokaryotic Transmembrane β-barrel Proteins. Journal of Molecular Biology, 433, 166729.

Mahrenholz, C.C. et al. (2011) Complex networks govern coiled-coil oligomerization--predicting and profiling by means of a machine learning approach. Mol Cell Proteomics, 10, M110.004994.

Manfredi, M. et al. (2022) E-SNPs&GO: embedding of protein sequence and function improves the annotation of human pathogenic variants. Bioinformatics, 38, 5168–5174.

Manfredi, M. et al. (2023) ISPRED-SEQ: Deep Neural Networks and Embeddings for Predicting Interaction Sites in Protein Sequences. Journal of Molecular Biology, 167963.

Rives, A. et al. (2021) Biological structure and function emerge from scaling unsupervised learning to 250 million protein sequences. Proc Natl Acad Sci U S A, 118, e2016239118.

Szczepaniak, K. et al. (2020) A library of coiled-coil domains: from regular bundles to peculiar twists. Bioinformatics, 36, 5368–5376.

Testa, O.D. et al. (2009) CC+: a relational database of coiled-coil structures. Nucleic Acids Res, 37, D315-322.

Trigg, J. et al. (2011) Multicoil2: predicting coiled coils and their oligomerization states from sequence in the twilight zone. PLoS One, 6, e23519.

Truebestein, L. and Leonard, T.A. (2016) Coiled-coils: The long and short of it. Bioessays, 38, 903–916.

Vincent, T.L. et al. (2013) LOGICOIL--multi-state prediction of coiled-coil oligomeric state. Bioinformatics, 29, 69–76.

Walshaw, J. and Woolfson, D.N. (2001) Socket: a program for identifying and analysing coiled-coil motifs within protein structures. J Mol Biol, 307, 1427–1450.

Wilson, I.A. et al. (1981) Structure of the haemagglutinin membrane glycoprotein of influenza virus at 3 A resolution. Nature, 289, 366–373.

Woolfson, D.N. (2023) Understanding a protein fold: The physics, chemistry, and biology of α-helical coiled coils. Journal of Biological Chemistry, 299, 104579.

Zemla, A. et al. (1999) A modified definition of Sov, a segment-based measure for protein secondary structure prediction assessment. Proteins: Structure, Function, and Bioinformatics, 34, 220–223.

